# A Perspective on Interaction Tests in Genetic Association Studies

**DOI:** 10.1101/019661

**Authors:** Hugues Aschard

**Affiliations:** Harvard School of Public Health, Department of Epidemiology, Boston, MA 02115, USA

## Abstract

The identification of gene-gene and gene-environment interaction in human traits and diseases is an active area of research that generates high expectation, and most often lead to high disappointment. This is partly explained by a misunderstanding of some of the inherent characteristics of interaction effects. Here, I untangle several theoretical aspects of standard regression-based interaction tests in genetic association studies. In particular, I discuss variables coding scheme, interpretation of effect estimate, power, and estimation of variance explained in regard of various hypothetical interaction patterns. I show first that the simplest biological interaction models—in which the magnitude of a genetic effect depends on a common exposure—are among the most difficult to identify. Then, I demonstrate the demerits of the current strategy to evaluate the contribution of interaction effects to the variance of quantitative outcomes and argue for the use of new approaches to overcome these issues. Finally I explore the advantages and limitations of multivariate models when testing for interaction between multiple SNPs and/or multiple exposures, using either a joint test, or a test of interaction based on risk score. Theoretical and simulated examples presented along the manuscript demonstrate that the application of these methods can provide a new perspective on the role of interaction in multifactorial traits.

## Introduction

Hundreds of studies have searched for gene-gene and gene-environment interaction effects in human data with the underlying motivation of identifying or at least accounting for potential biological interaction. So far, this quest has been quite unsuccessful and the large number of methods that have been developed to improve detection^1-5^ have not qualitatively changed this situation. This lack of discovery in the face of a large research investment has been discussed in several review papers that have pointed out a number of issues specific to interaction tests, including exposure assessment, time-dependent effect, confounding effect and multiple comparisons^2; 6; 7^. While these factors are obvious barriers to the identification of interaction effects, it appears that some of the limitations of standard regression-based interaction tests that pertain to the nature of interaction effects are greatly underestimated. Previous work showed the detection of some biologically meaningful interaction effects requires larger sample sizes than marginal effects for a similar effect size^8; 9^, however it is not an absolute rule. Understanding the theoretical basis of this lack of power can help us optimizing study design to improve detection of interaction effect in human traits and diseases, and open the path for new methods development. Moreover the interpretation of effect estimates from interaction models often suffer from various imprecisions. Compared to marginal models, the coding scheme for interacting variables can impact effect estimates and association signals for the main effects^9^. Also, the current strategy to derive the contribution of interaction effects to the variance of an outcome greatly disadvantages interaction effects and are inappropriate when the goal of a study is not prediction but to assess the relative importance of an interaction term from a biological perspective. While alternative approaches exist, they have not so far been considered in genetic association studies. Finally, the development of new pairwise gene-gene and gene-environment interaction tests is reaching some limits, because the number of assumption that can be leveraged to improve power is limited when only two predictors are considered. With the exponential increase of available genetic and non-genetic data, the development and application of multivariate interaction tests offer new opportunities to building powerful approaches and moving the field forward.

## Methods and Results

### Coding scheme and effect estimates

Consider an interaction effect between a single nucleotide polymorphism (SNP) *G* and an exposure *E* (which can be an environmental exposure or another genetic variant) on a quantitative outcome Y. For simplicity I assume in all further derivation that *E* is normally distributed with variance 1, and *G* and *E* are independents. The simplest and most commonly assumed underlying model for Y (i.e. the model used to generate the values of Y) when testing for an interaction effect between *G* and *E* is defined as follows:

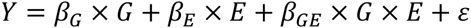

where *β*_*G*_ is the main effect of *G*, *β*_*E*_ is the main effect of *E*, *β*_GE_ is a linear interaction between *G* and *E*, and ε, the residual, is normally distributed with mean 0 and variance *σ*^2^ sets so that the variance of Y equals 1. One can then evaluate the impact of applying linear transformation of the genotype and/or the exposure when testing for main and interaction effects. For example, assuming *E* has a mean > 0 and *G* is defined as the number of coded allele in the generative model, *Y* can be rewritten as a function of *G*std and *E*std, the standardized *G* and *E*:

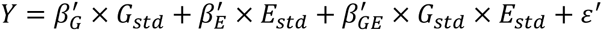

where 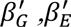 and 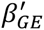 are the main effects of *G*_*std*_ and *E*_*std*_ and their interaction. Relating the standardized and unstandardized equations, we obtain (**Appendix A**):

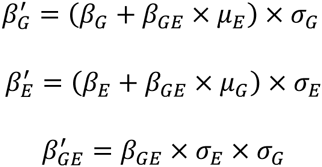

where μ_*G*_, *σ*_*G*_, μ_*E*_ and *σ*_*E*_ are the mean and variance of *G* and *E*, respectively. Hence, the estimated main effects of *G*_*std*_ and *E*_*std*_ not only scale with the variance of *G* and *E* but can also change qualitatively if there is an interaction effect. In comparison, the interaction effect 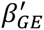 only scales with the predictors variance, however, because 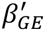 does not depend on *σ*_*GE*_ the variance of the interaction term but on the variance of *G* and *E*, the magnitude of the interaction effect can change.

Which coding scheme for *G* and *E* has the most biological sense can only be discussed on a case by case basis and is therefore out of the scope of this paper. The important point here, is that coding scheme should be chosen carefully when testing an interaction as it can correspond to profoundly different patterns on the outcome. This is illustrated in **Figure 1**, which shows the contribution of a pure interaction effect (*β*_*G*_ = *β*_*E*_ = 0 and β_*GE*_ ≠ 0) to Y. When *G* and *E* are centered, the joint effect of *G* and *E* is similar across the most extreme sub-groups (low exposure and homozygote for the protective allele *vs* high exposure and homozygote for the risk allele) and opposite effect otherwise (**Figure 1a**). Conversely, when *G* and *E* are positive or null, the interaction term simply corresponds to an increase (or decrease if the interaction term is negative) of the magnitude of a genetic effect when the exposure increases (**Figure 1b**). Hence, assuming *G* and/or *E* have a negative range in the generative model – besides it might have limited biological meaning – implies interaction effects of different nature as compared to model using positive predictors only. Furthermore, when the mean of the exposure increases while its variance is fixed, a realistic interaction effect for genetic data (i.e. explaining a small amount of the outcome variance) will appear more and more as a sole genetic effect (see **Figure S1**).

**Figure 1.**
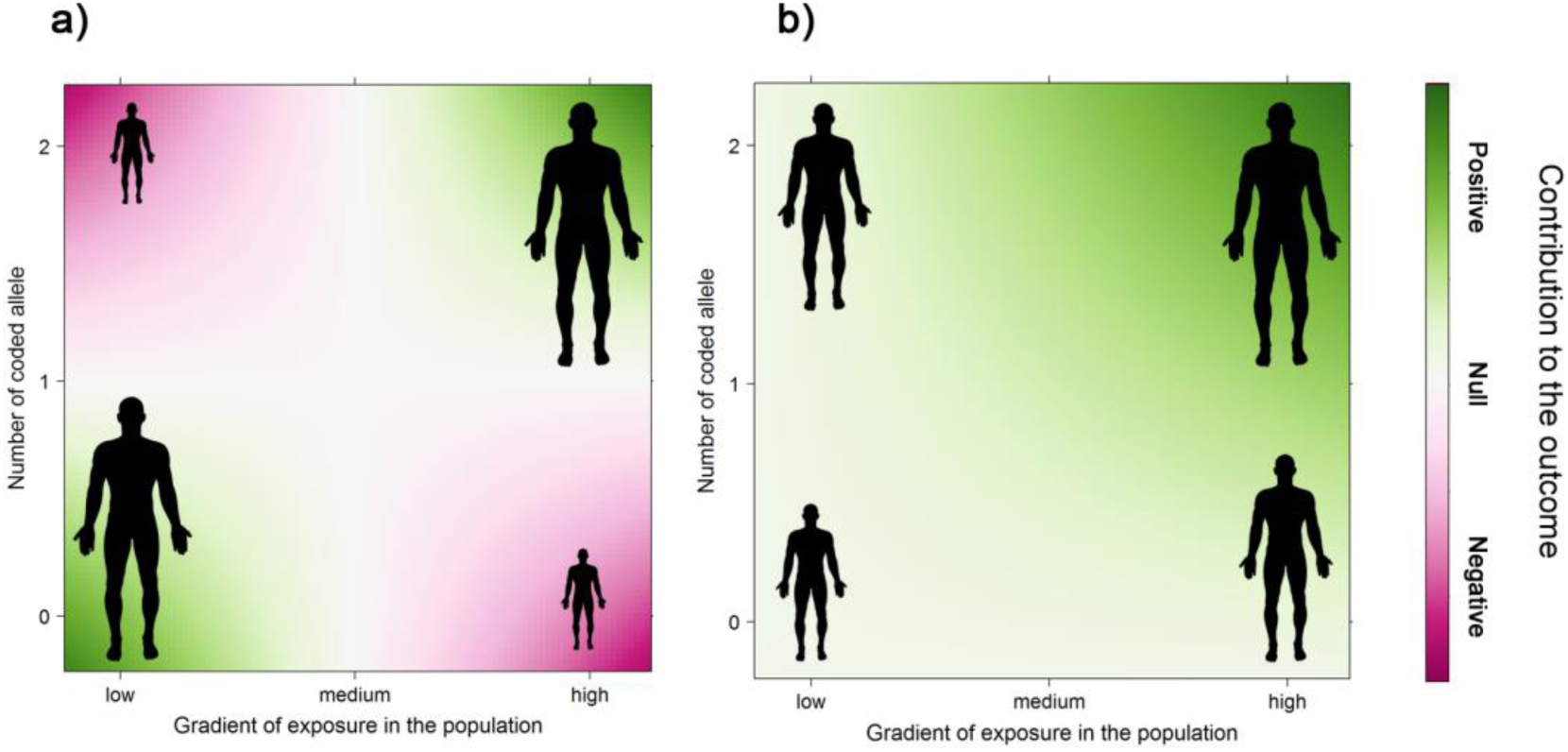
Examples of interaction patterns for a gene by exposure effect on height. Pattern of contribution of an interaction term to human height when shifting the location of the genetic variant and the exposure. In a) the interaction is defined as the product of centered genetic variant and exposure, while in b) genetic variant and exposure are positive or null.

While estimates for a specific coding scheme can be derived from estimates obtained from another coding scheme, questions arise on which final coding to choose and how to interpret estimates when modeling an interaction. This point, and the motivation for adding non-linear terms in general have been already debated and several general guidelines have been proposed (see for example the review by Robert J. Friedrich^10^). The consensus was that, if the range of the independent variables do naturally includes zero (e.g. smoking status, genetic variants) there is no problem in interpreting the estimated main and interaction effect. For an interaction effect between A and B, the main effect of A corresponds to the effect of A when B is null and conversely. Conversely, if the range of the variables do not naturally encompass zero, then the observed estimates “*will be an extrapolations beyond the observed range of experience"^10^.* Centering the variables can be an option to address this concern. In that case, the main effect of A and B would represent the effect of A among individuals having the mean value of B and conversely. However, as mentioned previously, using centered variables induces a less interpretable interaction term. A reasonable alternative consists in shifting the exposure values so that it has a minimum value close to 0, or alternatively to use ordinal categories of the exposure (e.g. high versus low BMI as done to define obesity), so that the main effect of A would correspond to the effect among the lowest observed value of B and conversely.

### Power considerations

The power of the tests from the interaction model and from a marginal genetic model defined as *Y* = *β*_*mG*_ × *G* + *ε*_*m*_, can be compared when deriving the non-centrality parameters (*ncp*) of the predictors of interest. Assuming all effects are small, so that *σ*^2^ the residual variance is close to 1, these *ncp* can be approximated by (see **Appendix B**):

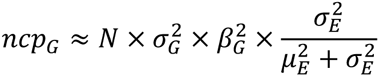

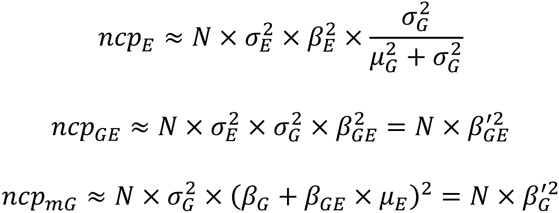

Note that in such scenario adjusting for the effect of *E* in the marginal genetic model has a minor impact on ncp_*mG*_. It would only be important in the presence of a strong exposure effect, such an effect would reduce *σ*^2^, the residual variance in the interaction model, and increase the *ncp*s from the interaction model but not ncp*mG*.

The above equations indicate first that the significance of the marginal test of *G* and the interaction test are invariant with the coding used in the model tested, while the significance of the test of the main genetic and exposure effects can change dramatically when shifting the mean of *G* and *E*. Second, as illustrated in **Figure 2,** depending on the parameters of the distribution of the exposure and the genetic variants in the generative model, the relative power of each test can be dramatically different. For example if the genetic variant has only a main linear effect but is not interacting with the exposure, we obtain 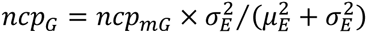, so that testing for *β*_*mG*_ will be much more powerful that testing for *β*_*G*_ if the mean of *E* is large, although there is no interaction effect here. When the generative model includes an interaction effect only (*β*_*G*_ = *β*_*E*_ = 0 and *β*_*GE*_ ≠ 0), we obtain 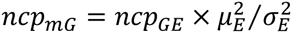. Again, the marginal test of the genetic effect can be dramatically more powerful than the test of interaction effect although the underlying model includes only an interaction term but no main effect.

**Figure 2.**
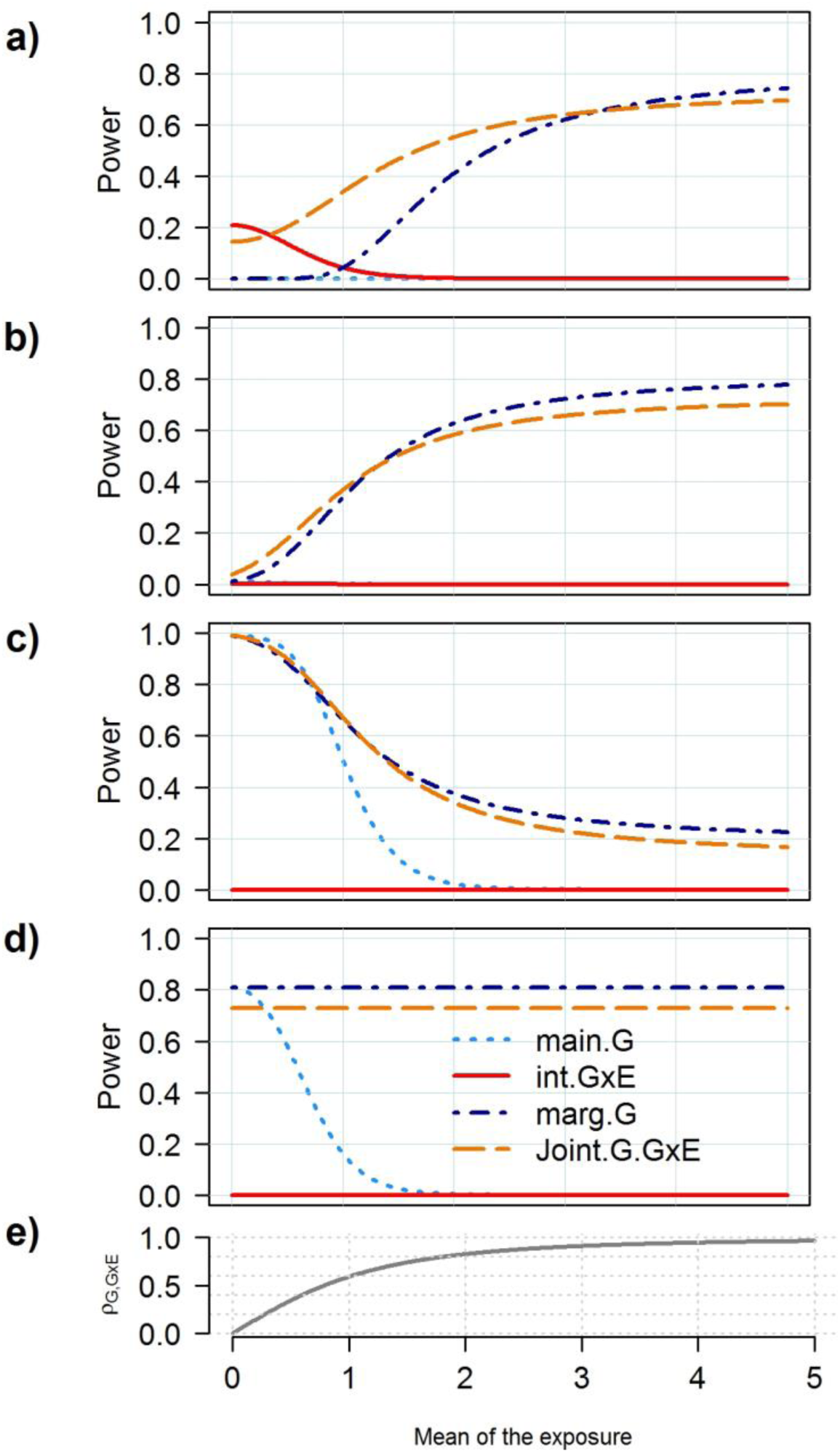
Relative power of the joint test of main genetic and interaction effects. Power comparison for the tests of the main genetic effect (*main.G*), the interaction effect (*int.GxE*) and the joint effect (*Joint G.GxE*) from the interaction model, and the test of the marginal genetic effect (*mar.G*). The outcome Y is define as a function of a genetic variant *G* coded as [0,1,2] with a minor allele frequency of 0.3, and the interaction of *G* with an exposure *E* normally distributed with variance 1 and mean 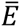 the genetic and interaction effects vary so that they explain 0% and 0.04% (a), 0.1% and 0.1% (b), 0.6% and 0.1% with effect in opposite direction, and 0.4% and 0% (d) of the variance of Y, respectively. Power and ρ*G*,*G*×*E*, the correlation between *G* and the *G* × *E* interaction term (e) were plotted for a sample size of 10,000 individuals and increasing 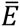 from 0 to 5.

More generally it follows that the power to detect an interaction effect explaining for example 1% of the variance of Y but inducing no marginal genetic effect (i.e. when *E* is centered as in **Figure 1a**) is much higher than for an interaction explaining the same amount of variance but whose effect can be capture by a marginal term (i.e. when *E* is not centered as in **Figure 1b-d**). This result is a direct consequence of the covariance between *β*_*G*_ and *β*_*GE*_ that arise when having non-centered exposure in the generative model (**Figure 2e**). This covariance equals 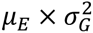 (**Appendix C**). It induces uncertainty on the estimation of the predictor effects, which decreases the significance estimates in the interaction model. With increasing inter-correlations between predictors it becomes impossible to disentangle the effects of one predictor from another, the standard errors of the effect estimates becoming infinitely large and the power decreases to the null^11^. As showed in the simulation study from **Figure S2**-**S3** these results are consistent for both linear and logistic regression and when assuming non-normal distribution of the exposure.

This lead to the non-intuitive situation where the power to detect a relatively simple and parsimonious interaction effect from a biological perspective – defined as the product of a genetic variant and an exposure both coded to be positive or null – is very small; and in most scenarios where the main genetic and interaction effects do not canceled each other (*see* e.g. ^12^) the marginal association test of *G* would be more powerful. In comparison a more exotic interaction effect as defined in **Figure 1a** and **Figure S1e**, would be both much easier to detect in a genome-wide interaction screening and not captured in a GWAS of marginal genetic effect.

### Proportion of variance explained

In genetic association studies the proportion of variance explained by an interaction term is commonly evaluated as the amount of variance of the outcome it can explain on top of the marginal linear effect of the interacting factors^13^. Following the aforementioned principle, one can derive the contribution of 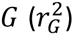, 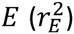 and 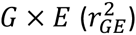to the variance of the outcome using the estimates from the standardize model, in which the interaction term is independent from *G* and *E*:

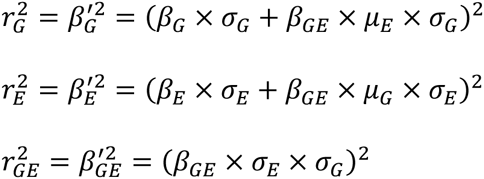

The total variance explained by the predictors in the interaction model equals 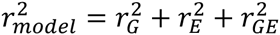(**Appendix D**). It follows that one can draw various scenarios where the estimated main effect of *E* and *G* can be equal to zero but have a non-zero contribution to the variance of Y because of the interaction effect. Indeed, the **Figure 3** shows that depending on the frequencies of the causal allele and the distribution of the exposure in the generative model, the vast majority of the contribution of the interaction term to the variance of Y will be attributed to either the genetic variant or the exposure. This is in agreement with recent work showing that even if a large proportion of the genetic effect on a given trait is induced by interaction effects, the observed contribution of interaction terms to the heritability can still be very small^13^. Because such interaction effects have small contribution to 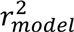on top of the marginal effects of *E* and *G*, they have a very limited utility for prediction purposes in the general population^14; 15^.

**Figure 3:**
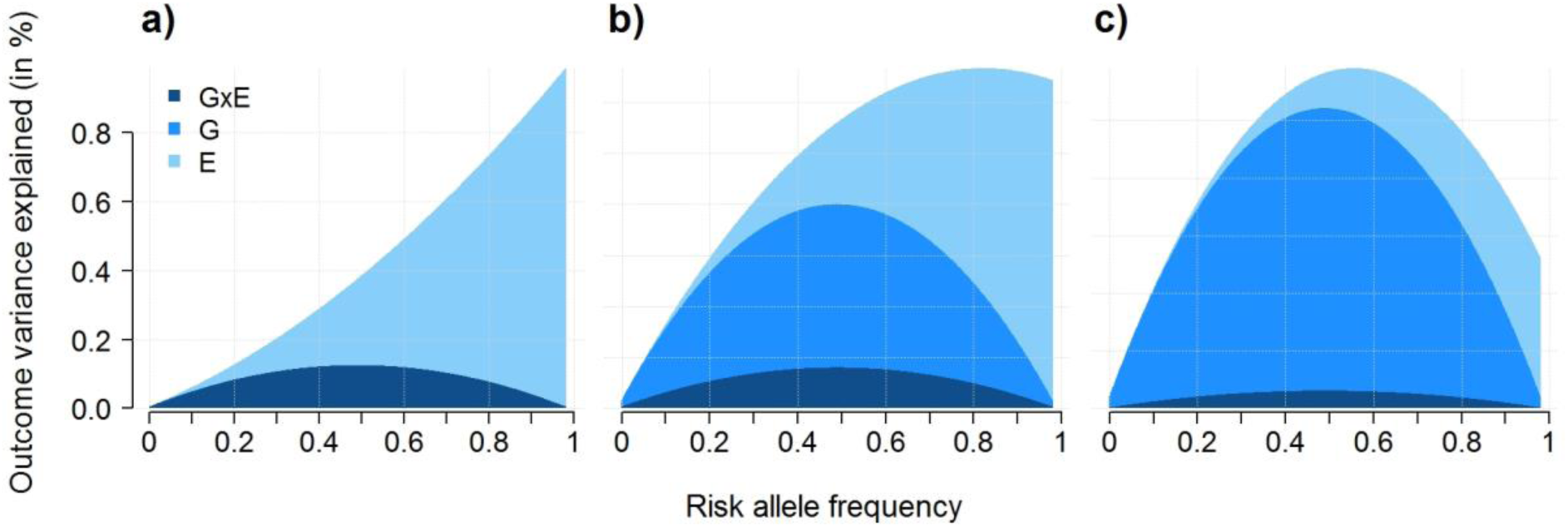
Examples of attribution of phenotypic variance explained by an interaction effect. Proportion of variance of an outcome *Y* explained by a genetic variant *G*, an exposure *E* and their interaction *G*x*E* in a model harboring a pure interaction effect only (Y = *β*_*GE*_ × *G* × *E* + ε). The exposure *E* follows a normal distribution with a standard deviation of 1 and mean of 0 (a), 2 (b) and 4 (c). The genetic variant is biallelic with a risk allele frequency increasing from 0.01 to 0.99. The interaction effect is set so that the maximum of the variance explained by the model equals 1%.

This is a strong limitation when the goal is not prediction but to understand the underlying architecture of the trait under study and to evaluate the relative importance of main and interaction effects from a public health perspective. Lewontin^16^ highlighted this issue a few decades ago, showing that the analysis of causes and the analysis of variance are not necessarily overlapping concepts. His work presents various scenarios where “*the analysis of variance will give a completely erroneous picture of the causative relations between genotype, environment, and phenotype because the particular distribution of genotypes and environments in a given population*". Since then, a number of theoretical studies have explored the issue of assigning importance to correlated predictors^17-20^ and several alternatives measures have been proposed. To my knowledge, none of these measures has been considered so far in human genetic association studies. The advantages and limitation of these alternatives have been debated for years and no clear consensus arose, however Pratt axiomatic justification^21^ for one of these measures – further presented in the literature as the Product Measure^22^, Pratt index or Pratt’s measure^23^ – has various interesting properties that makes it a relevant substitute. For a predictor X_i_, the Pratt’s index that we refer further as r^2^*, is defined as the product of *β*X_*i*_, the standardized coefficient from the multivariate model (where all predictors are scaled to have mean 0 and variance 1, including the interaction term), times its marginal (or zero-order) correlation with the outcome cor(*Y*, *X*_*i*_), i.e. 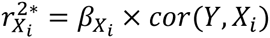.

By definition, 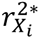 attributes a predictor’s importance as a direct function of its estimated effect and therefore addresses the concern previously raised. Among other relevant properties, it depends only on regression coefficients, multiple correlation and residual variance but not higher moments, and it does not change with (non-constant) linear transformation of predictors other than *X*_*i*_. It also has convenient additivity properties as it satisfies the condition 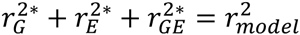 (**Appendix D**), so that the overall contribution of the predictors is the sum of their individual contribution, and for example the cumulated contribution of multiple interaction effects can easily be evaluated by summing 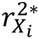. The Pratt’s index also received criticisms^20; 22^, in particular for allowing 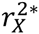 being negative^23^. Pratt’s 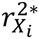 answer to this concern is that 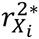 only describes the average contribution of a predictor to the outcome variance in one dimension and is therefore, as any one-dimension measure, a sub-optimal representation of the complexity of the underlying model. For example, a negative 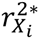 means that if we were able to remove the effect of *X*__i__, the variance of the outcome would increase because of the correlation of Xi with other predictors.

From a practical perspective, 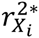 can be expressed as a function of the estimated effects, the means and the variances of *E* and *G* (**Appendix D**), and can therefore be derived from summary statistics of standard GWAS:

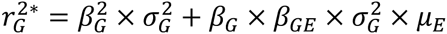

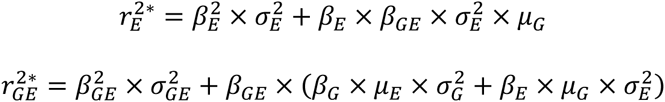

As showed in **Figure 4** and **Figure S4**, the Pratt index can recover the pattern of the causal model in situations where the standard approach would dramatically underestimate the contribution of the interaction effects. It can therefore be of great use in future studies to evaluate the importance of potentially modifiable exposures that influence the genetic component of multifactorial traits.

**Figure 4.**
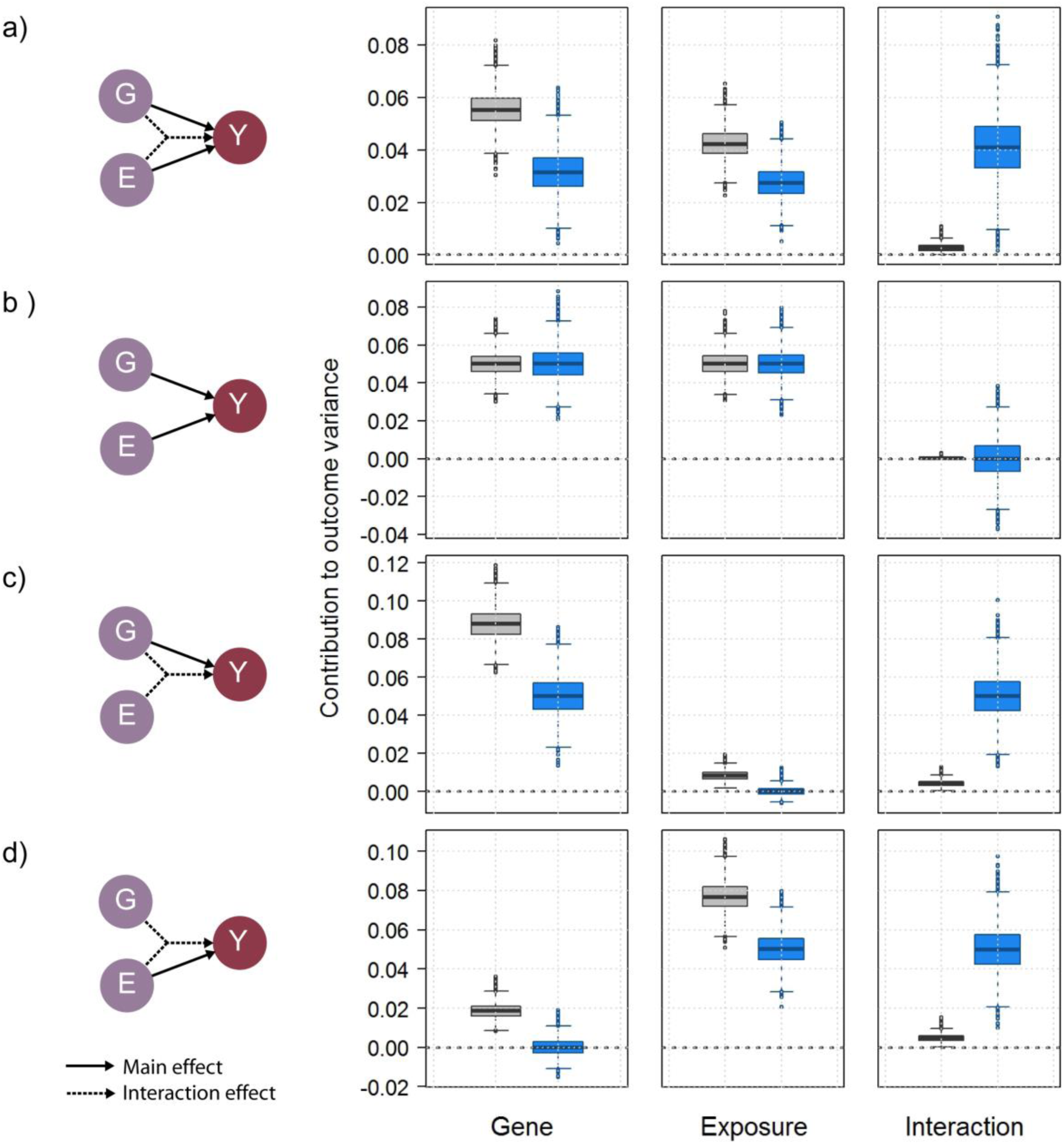
Importance of an interaction term as defined by the Pratt index. Contribution of a genetic variant *G* with minor allele frequency of 0.5, a normally distributed exposure *E* with mean of 4 and variance of 1 and their interaction *G*x*E,* to the variance of a normally distributed outcome *Y,* based on the standard approach – the marginal contribution of *E* and *G* and the increase in r^2^ when adding the interaction term– (grey boxes), and based on the Pratt index (blue boxes), across 10,000 replicates of 5,000 subjects. For illustration purposes the predictors explain jointly 10% of the variance of *Y*. In scenario a) all G, E and GxE have equals contribution, while in scenario b), c) and d) there was no interaction effect, no exposure effect, and no genetic effect, respectively.

### Improving detection through multivariate interaction tests

Using statistical technics such as the Pratt index can provide clues on the importance of interaction effects; however it does not help in mapping interaction. Increasing power mostly relies on two principles: increasing sample size, and leveraging assumptions on the underlying model. The case-only test, which assumes independence between the genetic variant and the exposure, and a two steps strategy which select candidate variants for interaction test based on their marginal linear effects or other parameters, are good examples of the later principle^4; 24; 25^. However, only a limited number of assumptions can be made for a single variant by a single exposure interaction test. With the overwhelming wave of genomic and environmental data, I suggest that a major path to move the field forward is to extend this principle while considering jointly more parameters.

This principle has first been applied over the past few years with the joint test of main genetic and interaction effects^26^. The *ncp* of such a joint test can be expressed as a function of main and interaction estimates (*β*_*G*_ and *β*_*GE*_), their variances (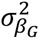 and 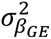) and their covariance *γ* (**Appendix E**).By accounting for *γ* the joint test recovers most of the power lost by the univariate test of the main genetic and interaction effect (so the situation where neither the interaction effect nor the main genetic effect are significant, while the joint test is, e.g. SNP rs11654749 in^27^). More importantly, in the presence of both main and interaction effects, it can outperform the marginal test of *G*. Although this is at the cost of decreased precision, i.e. if the test is significant, one cannot conclude whether association signal is driven by the main or the interaction effect. Moreover this would be true only if the contribution of the interaction effect on top of the marginal effect is large enough so that it balanced the increase in number of degree of freedom^28; 29^ (**Figure 2**).

Application of the joint test of main genetic effect and a single gene by exposure interaction term is now relatively common in GWAS setting^27; 30; 31^. However, exploring further multivariate interactions with multiple exposures is limited by practical considerations. Existing software to perform the joint test in a meta-analysis context^29; 32^ only allow the analysis of a single interaction term mostly because it requires the variance-covariance matrix between estimates, which is not provided by popular GWAS software. Leveraging the results from the previous sections on can show that the *ncp* of the joint test of main genetic effect and interactions with *l* independent exposures can be expressed as the sum of *ncp* from the test of *G* and the *G* × *E*_*cent.i*_ where *E*_*cent.i*_ is the centered exposure i (**Appendix E**):

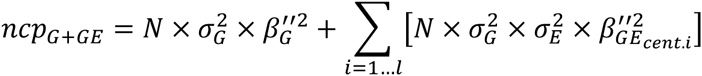

where 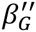 and 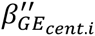 are the effects of *G* and *G* × *E*cent.i. Such a test is robust to non-normal distribution of the exposure, and realistic correlation (<0.1) between the genetic variant and the exposures, but sensitive to the relatively high correlation (>0.1) that could be observed between exposures (**Figure S5-S6**). Hence, one can perform meta-analysis of a joint test including multiple interaction effects using existing software simply by centering exposures. In brief one would have to perform first a standard inverse-variance meta-analysis to derive chi-squares for the l + 1 terms from the model considered, and then to sum all chi-squares to form a chi-square with l + 1 *df*. Importantly, centering the exposures will be of interest only when testing jointly multiple interactions and the main genetic effect. In comparison, the combined test of multiple interaction effects can be simply performed by summing chi-squares from each independent interaction test or from interaction test derived in a joint model. As previously, the validity of this approach relies on independence between the genetic variant and the exposures, and between the exposures. Finally, a more general solutions that should be explored in future studies would consists, as proposed for the analysis of multiple phenotypes (e.g.^33^), in estimating the correlation between all tests considered (main genetic effect and/or multiple interaction effects) using genome-wide summary statistics in order to form a multivariate test.

A second major direction for the development of multivariate test is to assume the effects of multiple genetic variants depend on a single “scaling” variable *E*. Various powerful tests can be built under such an assumption. A rising approach consists in testing for interaction between the scaling variable and a genetic risk score (GRS), derived as the weighted sum of the risk alleles. Several interaction effects have been identified using this strategy^34-39^, some being replicated in independent studies36; 37. This relative success, as compared to other univariate analysis, has generated discussion regarding potential underlying mechanisms^15; 40-43^. Overall, testing for an interaction effect between a GRS and a single exposure consists in expanding the principle of a joint test of multiple interactions while leveraging the assumption that, for a given choice of coded alleles, most interaction effects are going in the same direction. It is similar in essence to the burden test that has been widely used for rare variant analysis^44^. In its simplest form it can be expressed as the sum of all interaction effects and it captures therefore deviation of the mean of interaction effects from 0. When interaction effects are null on average, the joint test of all interaction tests (as previously described) will likely be the most powerful approach as it allows interaction effects to be heterogeneous. Conversely, if interactions tend to go in the same direction, the GRS-based test can outperform other approaches (**Figure 5)**. Of course, in a realistic scenario, a number of non-interacting SNPs would be included in the GRS, diluting the overall interaction signal and therefore decreasing power. However, the gain in power for the multivariate approaches can remain substantial even when a large proportion of the SNPs tested (e.g. 95% in the example from **Figure 5**) is not interacting with the exposure.

**Figure 5.**
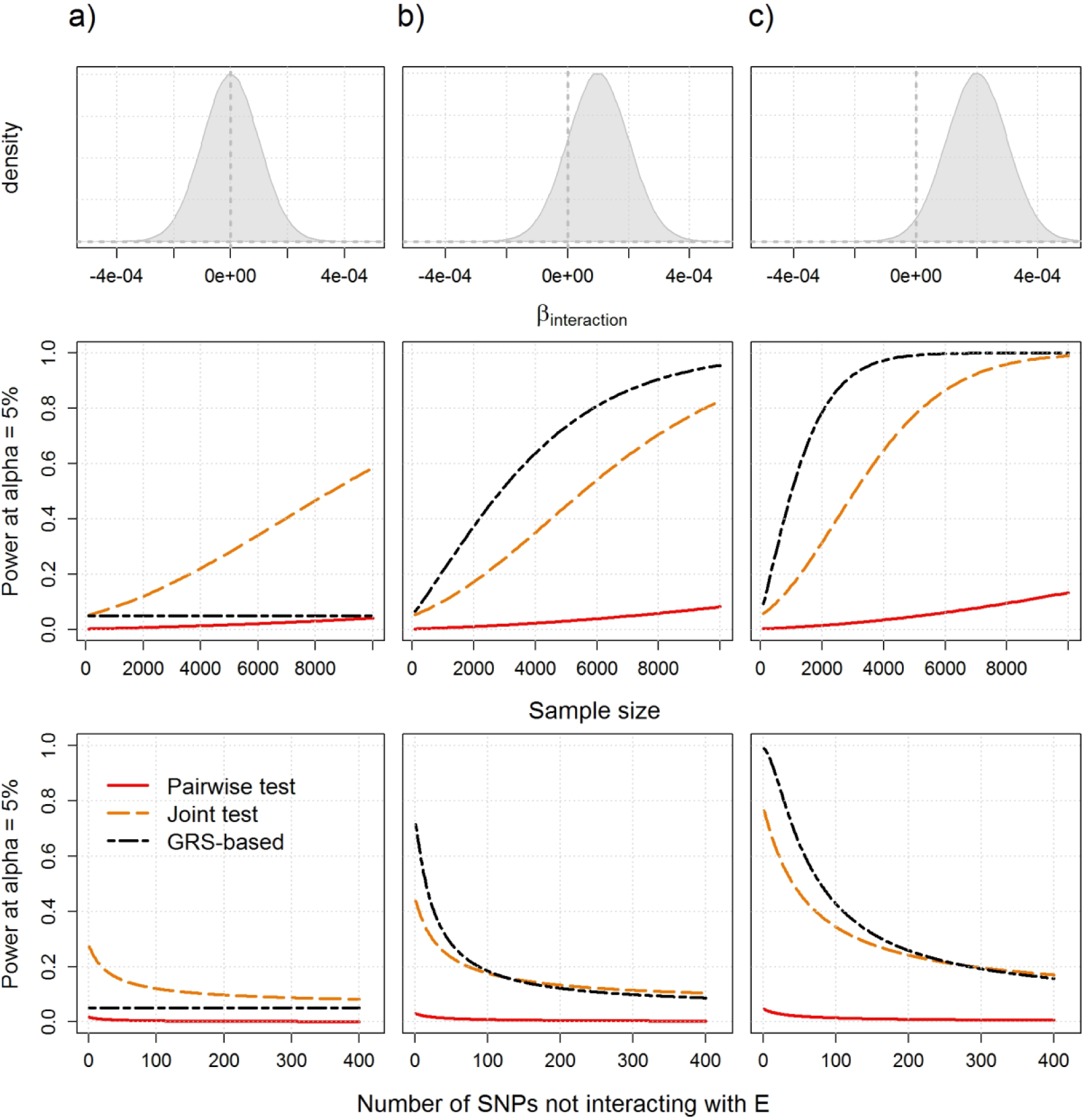
Advantages and limitation of testing interaction effect with a genetic risk score. Examples of power comparison for the combined analysis of interaction effects between 20 SNPs and a single exposure. Power was derived for three scenarios: the interaction effects are normally distributed (upper panels) and (a) centered, (b) slightly positive so that 25% of the interactions are negative, and (c) positive only. Three tests are compared while increasing sample size from 0 to 10,000: the joint test of all interaction terms, the genetic risk score by exposure interaction test, and the test of the strongest interaction effect after correction for the 20 tests performed (middle panels). The lower panels show power of the three tests for a sample size of 5,000, when including 1 to 400 non-interacting SNPs on top of the 20 causal SNPs in the analysis.

As showed in **Appendix F**, assuming the SNPs in the GRS are independents, the GRS by *E* interaction test can be derived from individual interaction effect estimates. More precisely, consider testing the effect of a weighted GRS on *Y*:

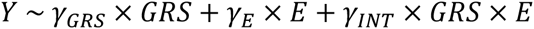

where *γ*_*GRSm*_, *γ*_*E*_ and *γ*_*INT*_ are the main effect of the weighted GRS, the main effect of *E* and the interaction effect between *E* and the GRS, respectively. The test of *γ*_*INT*_ is asymptotically equivalent to the meta-analysis of *γG*i×*E*, the interaction effects between *G*i and *E*, using an inverse-variance weighted sum to derive a 1 *df* chi-square, i.e. (see **Appendix F**, **Figure S7-S8,** and ^45^):

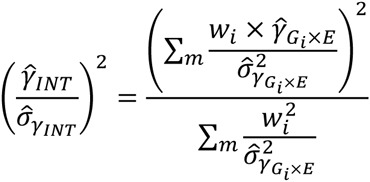

where *w*_*i*_ is the weight given to SNP i. A number of strategies can be used for the weighting scheme. In an agnostic search, assuming the interaction effects are independents of the SNPs characteristics, one should weight each SNP by the inverse of their standard deviation (*w*_*i*_ = 1/*σG*i). Alternatively, others have use weights proportional to the marginal genetic effect of the SNPs, assuming the magnitude of the marginal and interaction effects are correlated. The relative power of each of these weighting schemes depends on their relevance in regard to the true underlying model. Finally, applying GRS-based interaction tests implicitly supposed a set of candidate genetic variants have been identified. The current rationale consists in assuming that most interacting variants also display a marginal linear effect and therefore have focused on GWAS hits, however other screening methods can be used^46; 47^. Moreover existing knowledge, such as functional annotation^48^ or existing pathway database^49^ can be leverage to refine the sets of SNPs to be aggregated into a GRS.

## Discussion

Advancing knowledge of how genetic and environmental factors combine to influence human traits and diseases remains a key objective of research in human genetics. Ironically, the simplest and most parsimonious biological interaction models—those in which the effect of a genetic variant is either enhanced or decreased depending on a common exposure—are among the most difficult to identify. Furthermore, the contribution of such interaction effects can be dramatically underestimated when measured as the drop in *r*^2^ if the interaction term was removed from the model. Here, I argue for the use of new approaches and analytical strategies to address these concerns. This includes first using methods such as the Pratt index to evaluate the relative importance of interaction effects in genetic association studies. These methods can highlight important modifiable exposures influencing genetic mechanisms, which could be missed with the existing approach. Second, besides increasing sample size, increasing power to detect interaction effects in future studies will mostly rely on leveraging additional assumptions on the underlying model. In the big data era, where millions of genetic variants are measured on behalf of multiple environmental exposures and endo-phenotypes, this means using multivariate models. A variety of powerful statistical tests can be devised assuming multiple environmental exposures interact with multiple genetic variants. As showed in this study, the application of such approaches can dramatically improve power to detect interaction than standard univariate tests. While these methods comes at the cost of decreased precision—i.e. a significant signal would point out multiple potential culprit—they can identify interaction effects that would potentially be of greater clinical relevance that univariate pairwise interaction^14; 15^. The application of these methods in genetic association studies offers great opportunities for moving the field forward and providing a new perspective on interaction effects in human traits and diseases.

## Appendices

## Appendix A: Effect estimates from standardized and unstandardized predictors

Following the notation defined from the main text, the outcome *Y* can be expressed as:

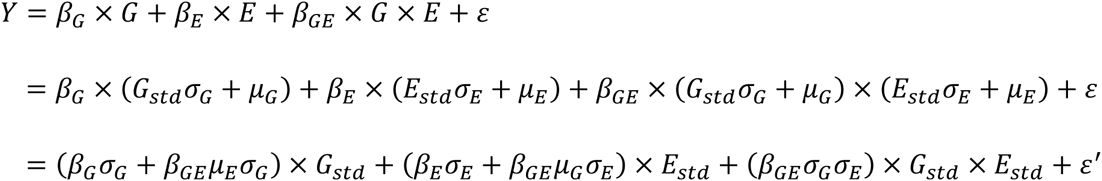

where *ε*’ is a term that depends neither on *G*_*std*_ nor *E*_*std*_. This leads to the following relationship between the standardized and unstandardized estimates:

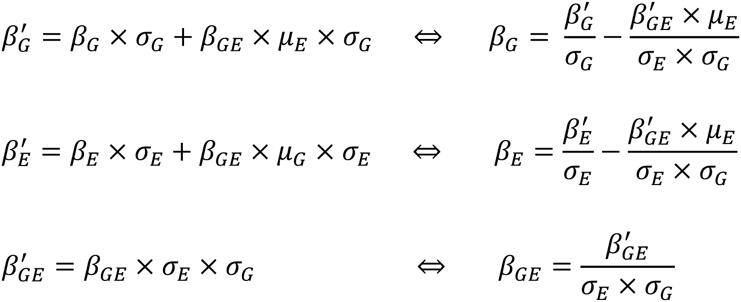

The variances of the unstandardized estimates equal:

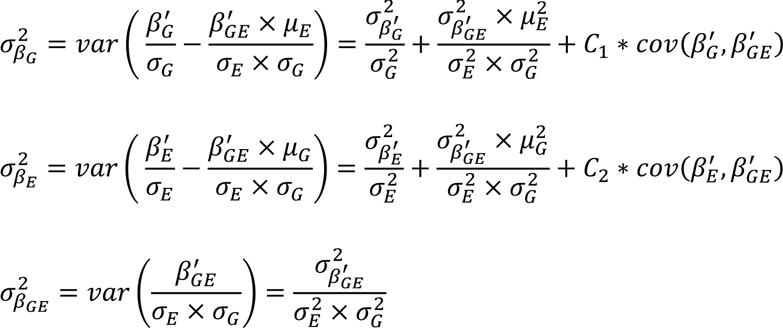

Where *C*_1_ and *C*_2_ are constants that depend on the mean and variance of *G* and *E*. **Appendix C** shows that 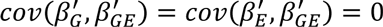 when *G* and *E* are independents. Moreover, when *G* and *E* are standardized, 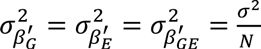, where *σ*^2^ is the residual variance of Y. Assuming the main effect of *G* and *E* and their interaction is small, so that *σ*^2^ *≈* 1, the variance of the estimates simplify:

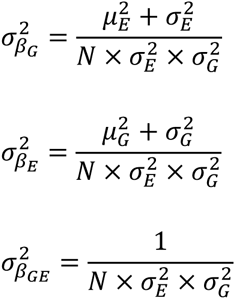

### Appendix B: Non-centrality parameters for marginal and interaction models

Using the estimates and variances from **Appendix A** one can derive *ncp*_*G*_, *ncp*_*E*_, and *ncp*_*GE*_, the non-centrality parameters (ncp) of the genetic main effect, the exposure main effect and the interaction effect under the assumptions of small effect sizes and *G* – *E* independence:

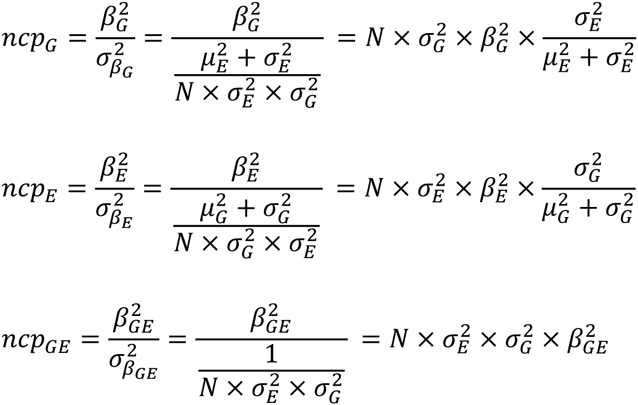

These *ncp* can be compared with ncp*mG*, the non-centrality parameter from the test of *G* in a marginal model. The marginal effect of *G*, *β*_*mG*_ is by definition the sum of the main effect of *G* plus the marginal contribution from interaction terms involving *G*. It can be approximate by:

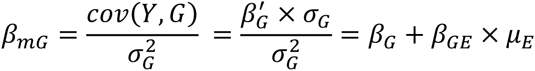

The marginal estimated effect of *E* can be derived similarly and equals:

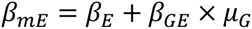

so that *ncp*_*mG*_ and *ncp*_*mE*_ (the *ncp* of the marginal test of *E*) can be expressed as follows:

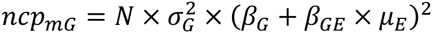

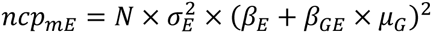

### Appendix C: Variance-covariance for the GxE term and its estimated effect

To derive the covariance and the correlation parameter between *G* and *G* × *E* we first calculate 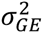, the variance of the interaction term *G* × *E* under the assumption of independence between *G* and *E*. Assuming the standard coding for *G* [0,1,2], and the frequency of the coded allele is ρ, and *E* is normally distributed so that *E*^2^ follows a non-central chi-square distribution with one degree of freedom, it can be express as:

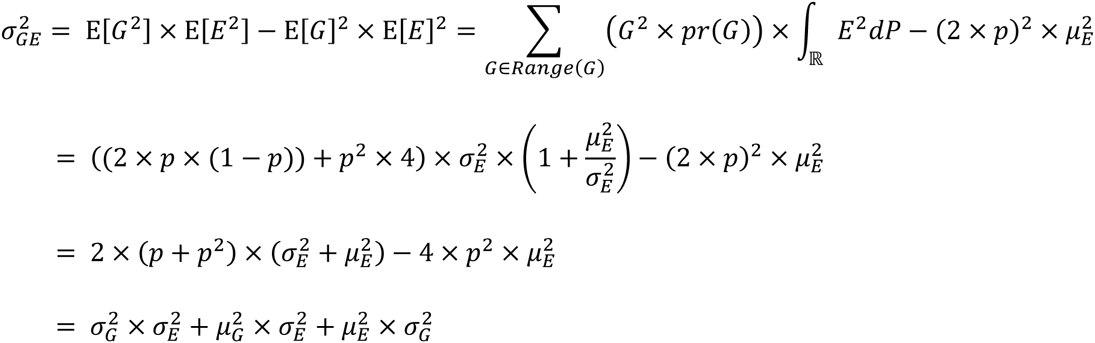

Under the same assumption, one can derive *cov*(*G*, *G* × *E*), the covariance between *G* and *G* × *E*:

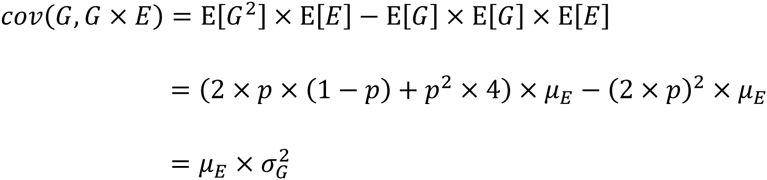

Similarly, one can derive the covariance between the exposure and the interaction and show that 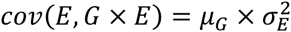. From this it appears that *cov*(*G*, *G* × *E*_*std*_) = *cov*(*E*, *G*_*std*_ × *E*) = *cov*(*G*_*std*_, *G*_*std*_ × *E*_*std*_)= 0.

The correlation between *G* and *G* × *E* equals then:

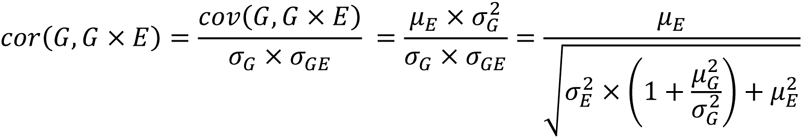

We derive then the covariance and correlation between the estimated effect of *G* and *G* × *E*. In its general form the variance-covariance matrix of estimates from the interaction model can be obtained using its matrix formulation: *σ*_*β*_ = (**X**^T^**X** –1*σ*^2^, where **X**, the matrix of predictor variables, equals [1, *G*, *E*, *G* × *E*] and *σ*^2^ is the variance of the residual of *Y*.

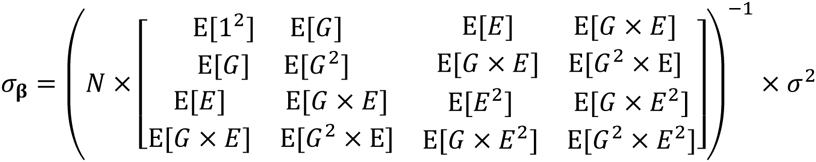

This is a relatively complex form, however when the predictors are standardized E[*E*] = E[*G*] = 0, and assuming *G* and *E* are independents, the formulation of *σ*_*β*_ greatly simplify, as all the off-diagonal elements of the matrix are null, so that:

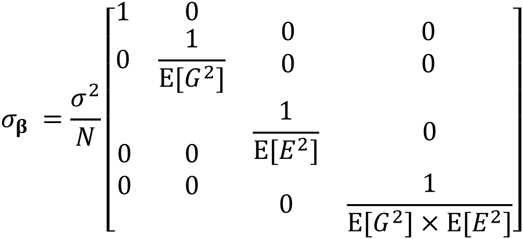

which implies that 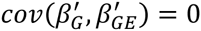. Building on this, and using the equations from **Appendix A**, we can derive *γ*, the covariance between *β*_*G*_ and *β*_*GE*_:

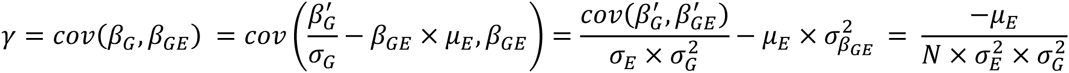

The correlation follows:

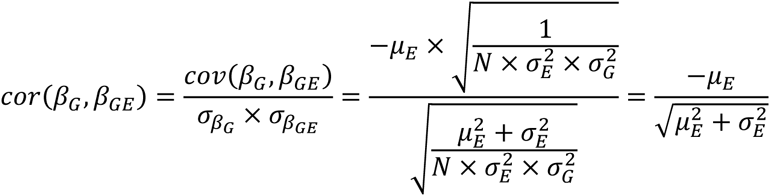

### Appendix D: Derivation of the Pratt index

To estimate the variance explained by predictors or other related measures, we first derive the expected variance of the outcome for a given generative model. For a single interaction term and assuming *G* – *E* independence, it equals:

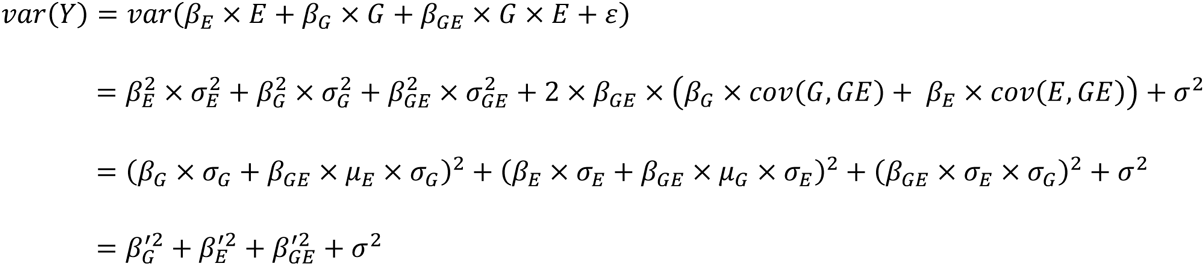

When more interaction terms are included, the outcome variance becomes a little more complex as additional covariance terms are added. For example assuming *k* interactions between *E* and *G*_*i*_, *i* = 1…*k*, the variance of *Y* becomes:

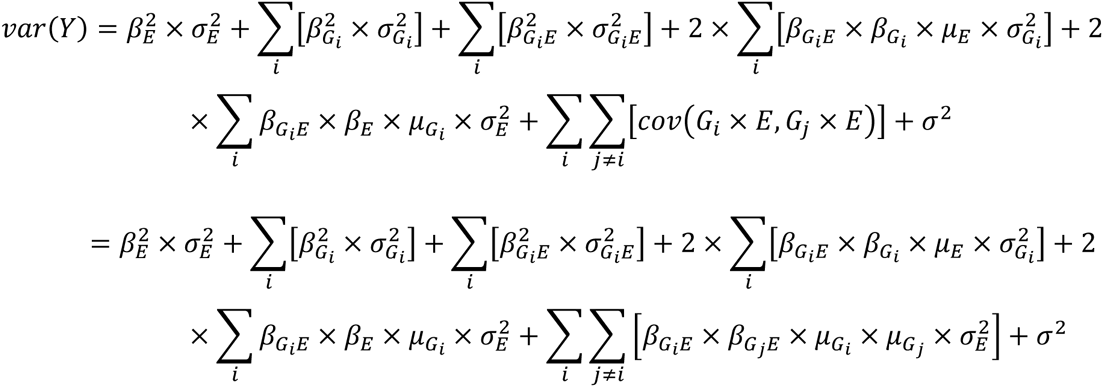

For simplicity let us assume *σ*^2^ is set so that *var*(*Y*) = 1 in all further derivation. When testing a single interaction term and using the equivalences from **Appendix A-B** one can show that the Pratt index can be expressed as a function of the estimates from the interaction model and the mean and variance of the genetic variant and the exposures considered:

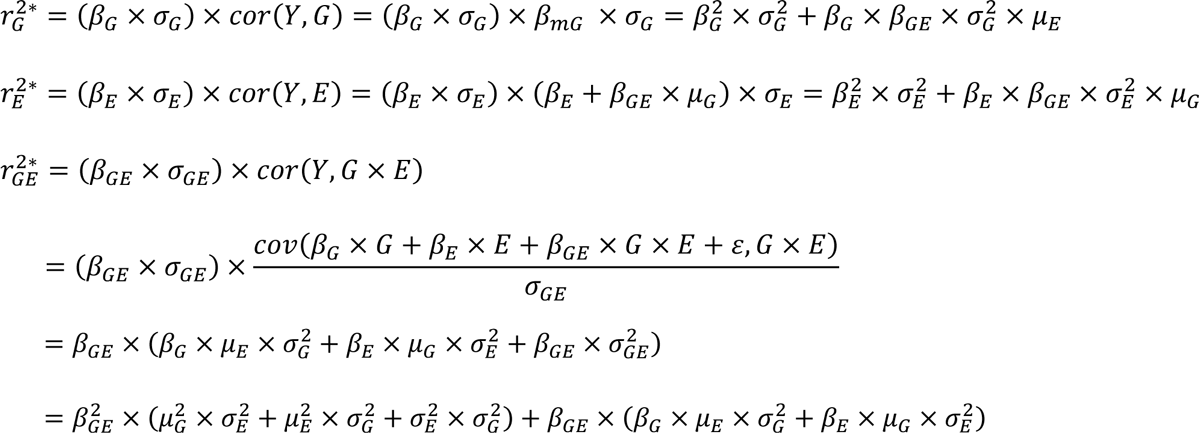

When summing the above Pratt index we obtain:

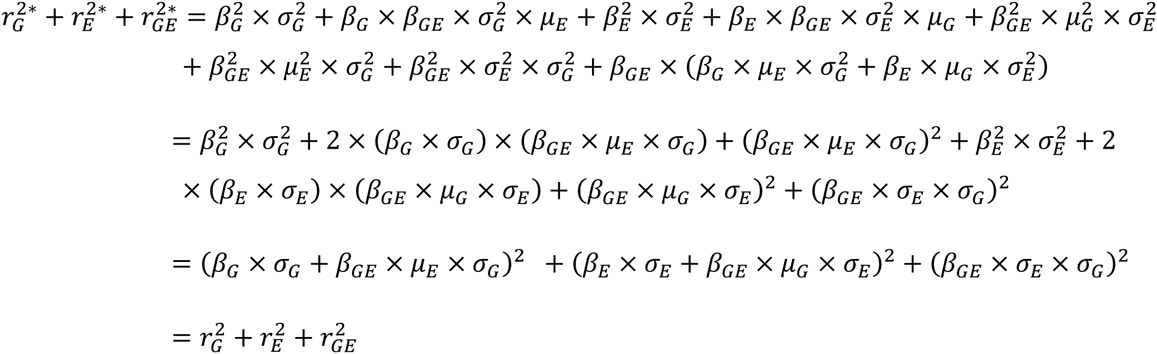

The cumulative contribution of multiple interactions involving independent SNPS can also be derived from summary statistics, although the derivation is a little less friendly because of additional covariance terms. For example assuming *k* interactions between *E* and *G*_i_, *i* = 1…*k*, we obtain (**Figure S4**) :

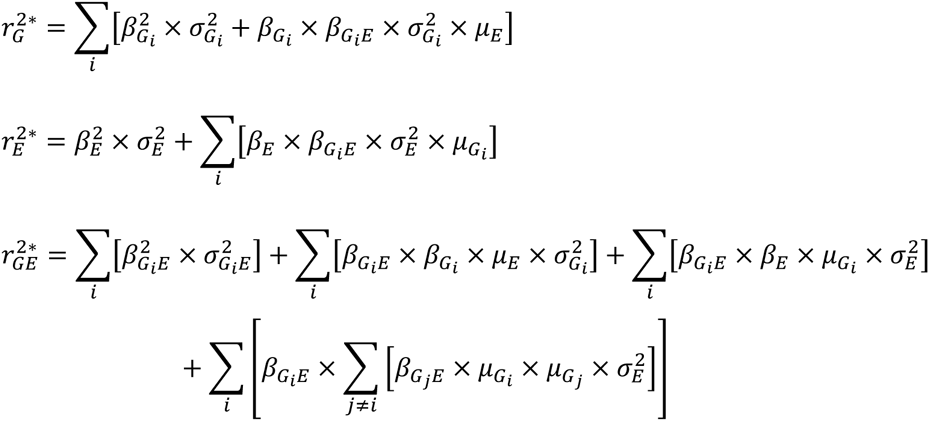

On should note that estimating the Pratt index for the exposure can be difficult in practice when the number of interaction is large, as it would require the estimated main exposure effect from a joint model including all SNPs main effect and all interactions term with the exposure. Also, because of the correlation between main and interaction terms, 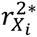, as the standard 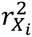, only approximate the amount the variance will change if *X*_*i*_ was held constant. For the latter measure, one can refer to^21^.

### Appendix E: Joint test of main and interaction effects

The multiple regression least square provides the estimated effect of the genetic main effect and interaction effects β = (*β*_*G*_, *β*_*GE*_) and their variance-covariance matrix ∑. The multivariate Wald test of the two parameters, which follow a 2 *df* chi-square can be expressed as:

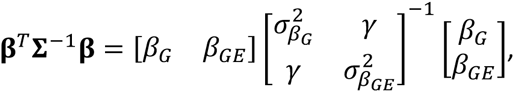

where *γ* is the covariance between *β*_*G*_ and *β*_*GE*_. It can be further developed as:

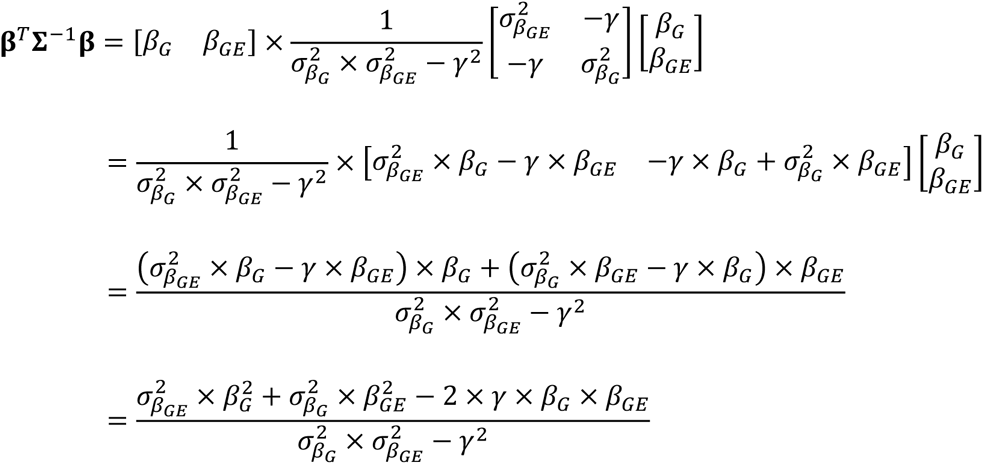

For clarity we derived the nominator and the denominator separately, so that β^*T*^*∑*–^1^β = *A*/*B*

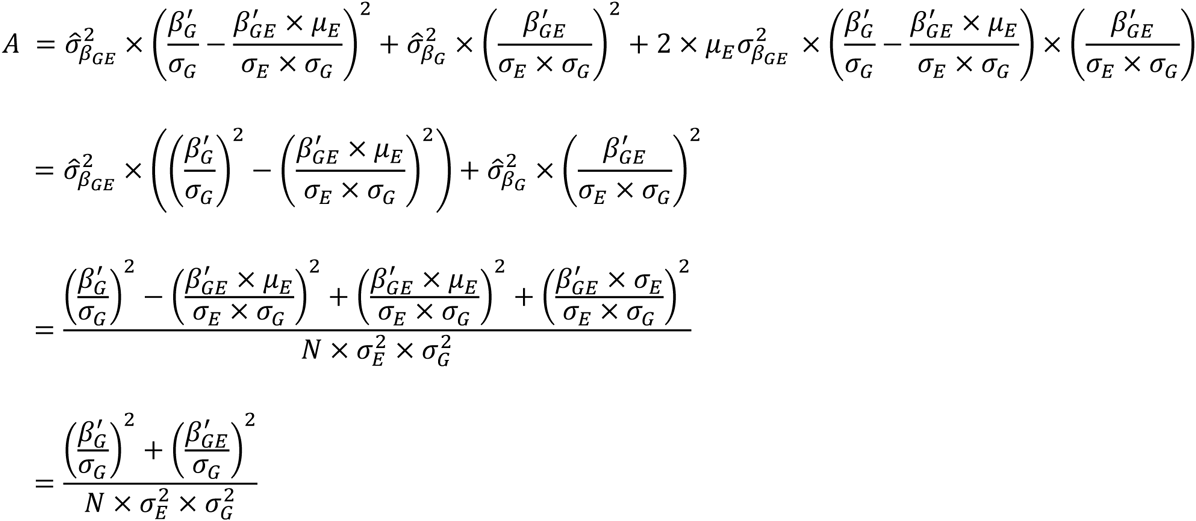

The denominator *B* equals:

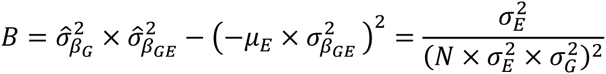

So that the joint test of *G* and *G* × *E* effects equals:

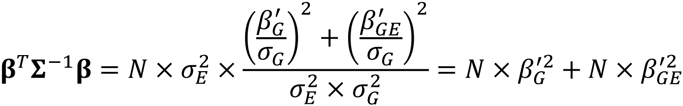

which is the sum of the individuals Wald test for the main effect and the interaction effect when *G* and *E* are standardized. Moreover, leveraging previous equivalences, we can express the joint test as a function of 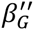 and 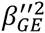, the estimated main and interaction effects from the model where *E* has been centered, so the test can be further expressed as:

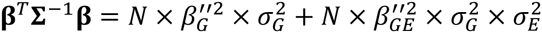

### Appendix F: GRS-based test, joint test and univariate test of multiple interaction effects

We denote β = (*β*_*G*1_, *β*_*G2*_,…*β*_*Gm*_) a vector of effects from *m* independent SNP, and 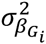and *w*_*i*_ are the variance of each estimate and weight of each SNP i in the genetic risk score (GRS), respectively. The effect of the weighted GRS on the outcome, *γG*RS, equals:

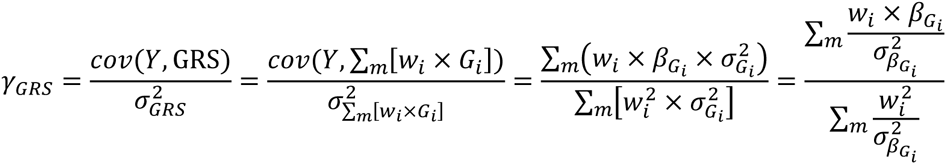

Consecutively, *σ*_*γGRS*_ the variance of *γ*_*GRS*_ can be derived as follows:

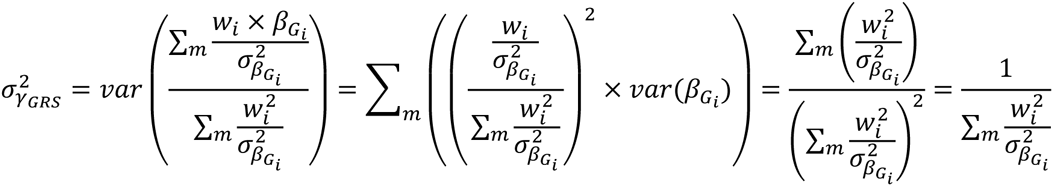

So that the chi-square of the marginal effect of the test of GRS equals:

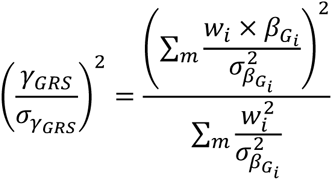

which corresponds to the inverse-variance weighted sum meta-analysis of each individual genetic variant. Similarly, one can derive the expected chi-square of the GRS by exposure interaction effect using *γ*_*Gi*×*E*_, the interaction effect between each SNP i and the exposure. Under the assumption of independence of the *m* interaction terms we obtain:

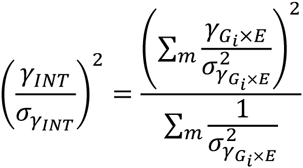

Hence for standardized *G* and *E* the *ncp* of the *GRS* by *E* interaction test equals 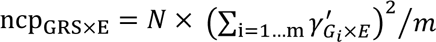 where *N* is the sample size and 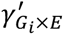 is the interaction effect from the standardized model, and follows a chi-square with one degree of freedom. In comparison, the *ncp* for the test of the strongest pairwise interaction, i.e. the interaction that explained the largest amount of variance, equals 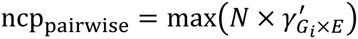.

## Supplemental Data

Supplemental Data include eight figures.

## Acknowledgements

I am grateful to Peter Kraft, Noah Zaitlen, Ami Joshi, John Pratt and Donald Halstead for helpful discussions and comments. This research was funded by NIH grants R03 HG006720.

## Web Resources

All simulations were conducted using the R sofware. http://www.r-project.org/

